# Early-childhood inflammation blunts the transcriptional maturation of cerebellar neurons

**DOI:** 10.1101/2022.07.26.501598

**Authors:** Seth A. Ament, Marcia Cortes-Gutierrez, Brian R. Herb, Evelina Mocci, Carlo Colantuoni, Margaret M. McCarthy

## Abstract

Inflammation early in life is a clinically established risk factor for autism spectrum disorders and schizophrenia, yet the impact of inflammation on human brain development is poorly understood. The cerebellum undergoes protracted postnatal maturation, making it especially susceptible to perturbations contributing to risk of neurodevelopmental disorders. Here, using single-cell genomics, we characterize the postnatal development of cerebellar neurons and glia in 1-5-year-old children, comparing those who died while experiencing inflammation vs. non-inflamed controls. Our analyses reveal that inflammation and postnatal maturation are associated with extensive, overlapping transcriptional changes primarily in two subtypes of inhibitory neurons: Purkinje neurons and Golgi neurons. Immunohistochemical analysis of a subset of these brains revealed no change to Purkinje neuron soma size but evidence for increased activation of microglia in those subjects experiencing inflammation. Maturation- and inflammation-associated genes were strongly enriched for those implicated in neurodevelopmental disorders. A gene regulatory network model integrating cell type-specific gene expression and chromatin accessibility identified seven temporally specific gene networks in Purkinje neurons and suggested that the effects of inflammation correspond to blunted cellular maturation.

**One Sentence Summary:** Post-mortem cerebelli from children who perished under conditions that included inflammation exhibit transcriptomic changes consistent with blunted maturation of Purkinje neurons compared to those who succumbed to sudden accidental death.

## INTRODUCTION

Early life inflammation is a clinically established risk factor for neurodevelopmental disorders, including autism spectrum disorders and schizophrenia (*1*). Animal models of maternal immune activation are providing insights into the cellular mechanisms of dysregulation (*2, 3*), but emphasis is focused on inflammatory insults that occur prenatally and their impacts in cortical regions. Adverse experiences that occur postnatally in early childhood, including inflammation, can also impact risk for neurodevelopmental disorders, yet the underlying mechanisms remain poorly understood.

The cerebellum -- an integral brain region not only for motor control but also for an array of higher cognitive functions including language processing, sociability, and emotionality (*4–7*) -- is one of the first brain regions to emerge during development and one of the last to reach maturity. Extensive cerebellar development occurs postnatally, and in humans the overall volume peaks in the early teens, occurring several years earlier in girls than boys, with both following an inverted U-shaped curve (*8*). The neural circuit in the cerebellum is comprised of repeating units consisting of a Purkinje neuron and its single climbing fiber from the inferior olivary nucleus, parallel fibers from granule neurons, and axonal output to the deep cerebellar nuclei (*9*). Molecular layer interneurons receive excitatory input from parallel fibers and Purkinje neurons (*10*). Golgi neurons establish extensive arborizations throughout the granule layer, receive excitatory input from both mossy fibers and granule cells, and provide recurrent inhibitory feedback to granule neurons (*11*). Imposed upon these units is a complex organization of parasagittal stripes and transverse zones, which create functional modules that greatly enhance the computational power and diverse functions of the cerebellum (*12, 13*).

The developmental generation of distinctly functional modules begins with early differentiation of Purkinje neurons into discrete populations by intrinsic genetic programs, a process nearing completion by the end of the first year of life in humans (*7*) and which subsequently establishes and constrains synaptic integration with other cellular components (*12*). Early on, each Purkinje neuron is innervated by multiple climbing fibers and by a process of activity dependent competition reduces to one for proper function, a process largely completed during the 2^nd^ postnatal week in rodents and early childhood in humans (*12, 14*). The complex orchestration of intrinsic genetic programs within Purkinje neurons and coordinated integration of multiple cellular inputs creates a moving target of sensitive and critical periods. When combined with relative immaturity at birth and an extensive period of postnatal maturation, the cerebellum becomes particularly susceptible to perturbation by extrinsic factors such as hypoxia/ischemia, toxins and inflammation (*7*). Abnormalities of the cerebellum are strongly associated with neurodevelopmental disorders including autism spectrum disorders, schizophrenia and attention and hyperactivity disorders (*15–22*).

Previously, we discovered a hidden critical period in cerebellar development in the early postnatal rat during which Purkinje neurons are sensitive to inflammation, with damage mediated by prostaglandin induced estradiol synthesis via stimulation of the aromatase enzyme (*23–25*). Preliminary evidence suggests a similar postnatal critical period in 1-9 year old human children, in whose cerebelli we detected elevated transcription of components of the same pathway in those that perished with conditions involving inflammation as opposed to sudden death due to accident (*26*). The human cerebellum remains under-represented in transcriptomic studies, and this is particularly true for its development (*27*).

Here we present a comprehensive analysis of single-cell transcriptomic profiles in the cerebellum in a sample of children aged 1-5 years with and without inflammation, as well as an extended developmental profile, integrating newly generated single-cell transcriptomic and epigenomic data from the adult human cerebellum and a recently described atlas for its prenatal development (*27*). Cell specific patterns of differential gene expression in cerebelli of children with inflammation were remarkably consistent and suggestive of blunted maturation based on the developmental profile as well as profound down regulation of many genes previously implicated by loss-of-function mutations as influencing risk for neurodevelopmental disorders.

## RESULTS

### A single-cell genomic atlas of the human cerebellum during its childhood maturation

We performed 10x Genomics single-nucleus RNA sequencing (snRNA-seq) of the cerebellar vermis or paravermis from 17 post-mortem human donors who were one-to five-years-old at the time of death (Table S1). Clustering of 168,719 high-quality cells revealed 19 major cell groups. We assigned these to known cerebellar cell types using established markers from previous studies in mice (*28*) (Fig. 1A,B; Table S2). For comparison, we also generated snRNA-seq of 51,388 cells from the post-mortem cerebellum of two adult human donors (Fig. 1C; Table S3). Marker gene correlations revealed a high degree of reproducibility for the cell types and their top markers in the cerebellum of children vs. adults (Fig. 1D).

**Figure 1.**
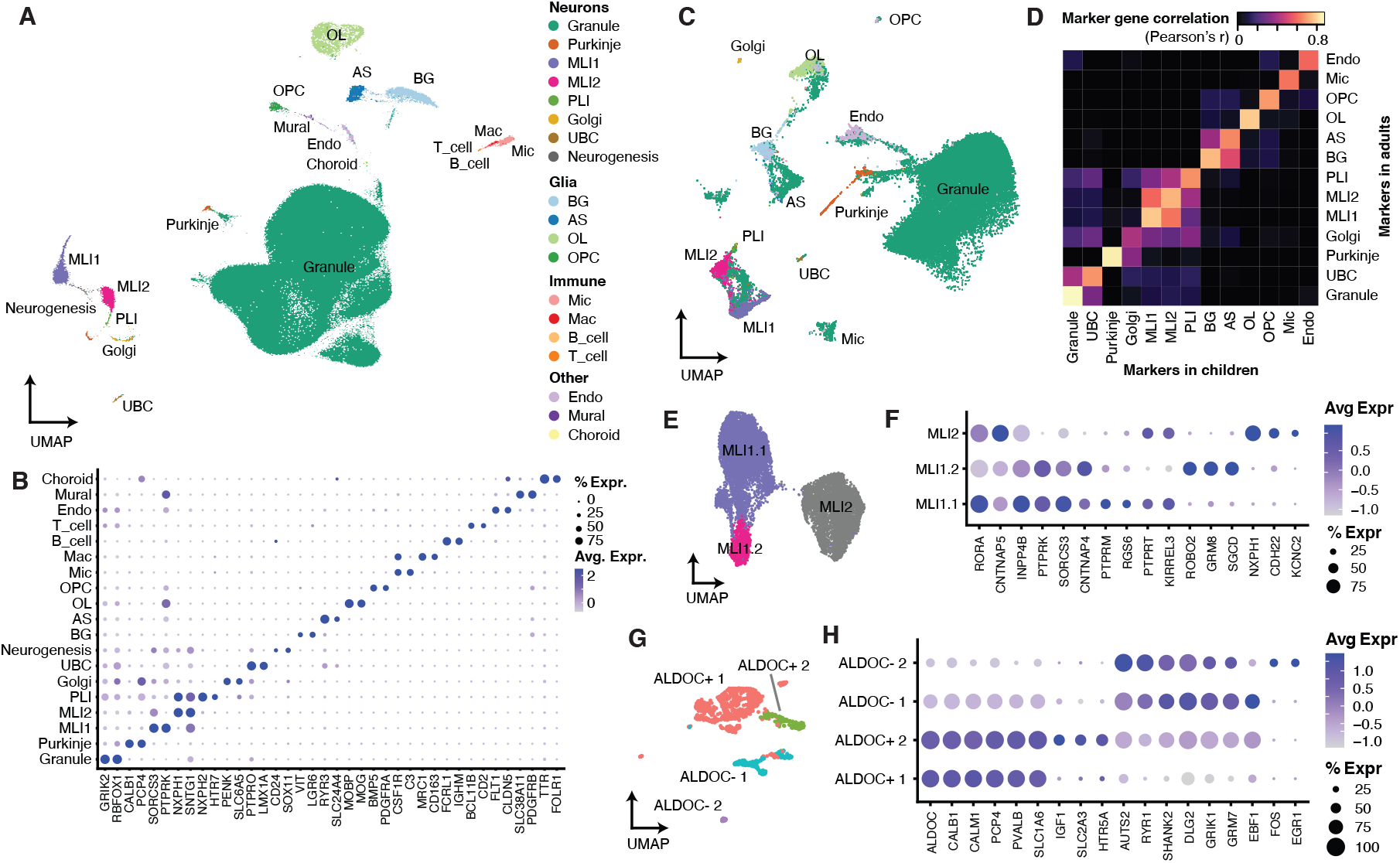
A single-cell genomic atlas of cell types in the human cerebellum. A. Uniform manifold approximation and projection (UMAP) for snRNA-seq of post-mortem human cerebellum from 1- to 5-year-old children. B. Expression of top cell type markers in snRN-seq of children. C. UMAP for snRNA-seq of the adult cerebellum. D. Correlation of cell type marker genes in the cerebellum of children vs. adults. E,F. Sub-clustering of molecular layer interneurons. G,H. Sub-clustering of Purkinje neurons. A web-based tool for the visualization of these snRNA-seq data is available at Neuroscience Multi-Omic Analytics (https://nemoanalytics.org/p?s=522872b1). MLI = molecular layer interneuron; PLI = Purkinje layer interneuron; UBC = unipolar brush cell; Mic = microglia; Mac = macrophage.

Sub-clustering revealed transcriptionally distinct subtypes for several classes of neurons. We found two discrete subtypes of molecular layer interneurons (MLI1 and MLI2, Fig. 1E). The top markers of MLI1 included *INPP4B, PTRPK,* and *SORCS3,* and the top markers for MLI2 included *NXPH1* and *CDH22* (Fig. 1F; Table S4). Equivalently named MLI subtypes with similar markers were characterized recently in the cerebellum of mice, where it was shown that the two subtypes are physiologically distinct but do not correspond to previously described morphological MLI subtypes (*28*). Similar proportions of MLI1 vs. MLI2 were detected in the cerebelli of adults vs. children, suggesting that these subtypes are not specific to a particular developmental stage.

Sub-clustering also revealed four sub-populations of Purkinje neurons. These were primarily distinguished by expression of *ALDOC* (also known as Zebrin-2), a canonical marker of parasagittal stripes (Fig. 1G). *ALDOC+* Purkinje neurons were further characterized by elevated expression of several canonical Purkinje neuron markers, including *CALB1, PCP4, PVALB,* and *SLC1A6* (also known as EAAT4), while *ALDOC-* Purkinje neurons had higher expression for markers such as *AUTS2, SHANK2, DLG2,* and *GRIK1* (Fig. 1H; Table S5). These results extend recent observations in mice demonstrating substantial transcriptional and functional differences among Purkinje neuron sub-populations (*13, 28*).

### Cerebellar cell type-specific transcriptomic consequences of brain inflammation

Samples were grouped into either inflamed (n=8) or non-inflamed (n=9). Sources of inflammation included viral and bacterial infections, as well as asthma, and in some cases the primary indicator of inflammation was the use of anti-inflammatory medications around the time of death (Table S1). Quantitative PCR of bulk cerebellar tissues from inflamed donors confirmed elevated levels of *TLR4* and *PTGS2* (also known as *COX-2*), reliable markers of inflammation (*26*). Non-inflamed donors primarily had sudden deaths due to accidents. None of the donors had a prior diagnosis with a neurodevelopmental disorder. There were no systematic differences between the two groups of subjects in variables such as age, gender, post-mortem interval, or race/ethnicity.

First, we examined neuroimmune cells to confirm classical signs of reactivity. Subclustering revealed four sub-populations of microglia, whose markers correspond to homeostatic (Mic1-2; *CX3CR1*+), phagocytic (Mic3; *SPP1*+), and pro-inflammatory (Mic4; *IL1B* and *CD83*+) subtypes (Fig. 2A,B; Table S6). Strikingly, Mic4 putatively pro-inflammatory microglia were 3.4-fold enriched in inflamed vs. non-inflamed donor brains (generalized linear model on proportions; 24% vs. 7% of microglia, *P* = 9.3e-15; Fig. 2C), Mic1 and Mic3 were correspondingly less abundant in inflamed brains. Thus, inflammation is associated with a shift toward microglial sub-populations expressing classical markers of reactive, pro-inflammatory states. We performed immunohistochemistry experiments using paraffin-embedded sections that were available from a subset of these inflamed and non-inflamed cerebelli (n=3-4 per group) to determine whether inflammation was also associated with morphological changes. Visualization of microglia by staining for AIF1, also known as IBA1 (Fig. 2D-E), indicated that microglia in inflamed cerebelli were more compact in morphology, with less ramified and thicker processes compared to controls, along with evident increased intensity of IBA1 labeling. These changes are consistent with pro-inflammatory, reactive microglial states (*1, 29, 30*). Quantification of these images confirmed a significant increase in the area covered by microglial processes, consistent with their amoeboid appearance, when compared to non-inflamed subjects (*p* < 0.001; Fig. 2F). These results confirm transcriptional and morphological signatures of immune cell reactivity in the cerebelli of donors who died while experiencing inflammation.

**Figure 2.**
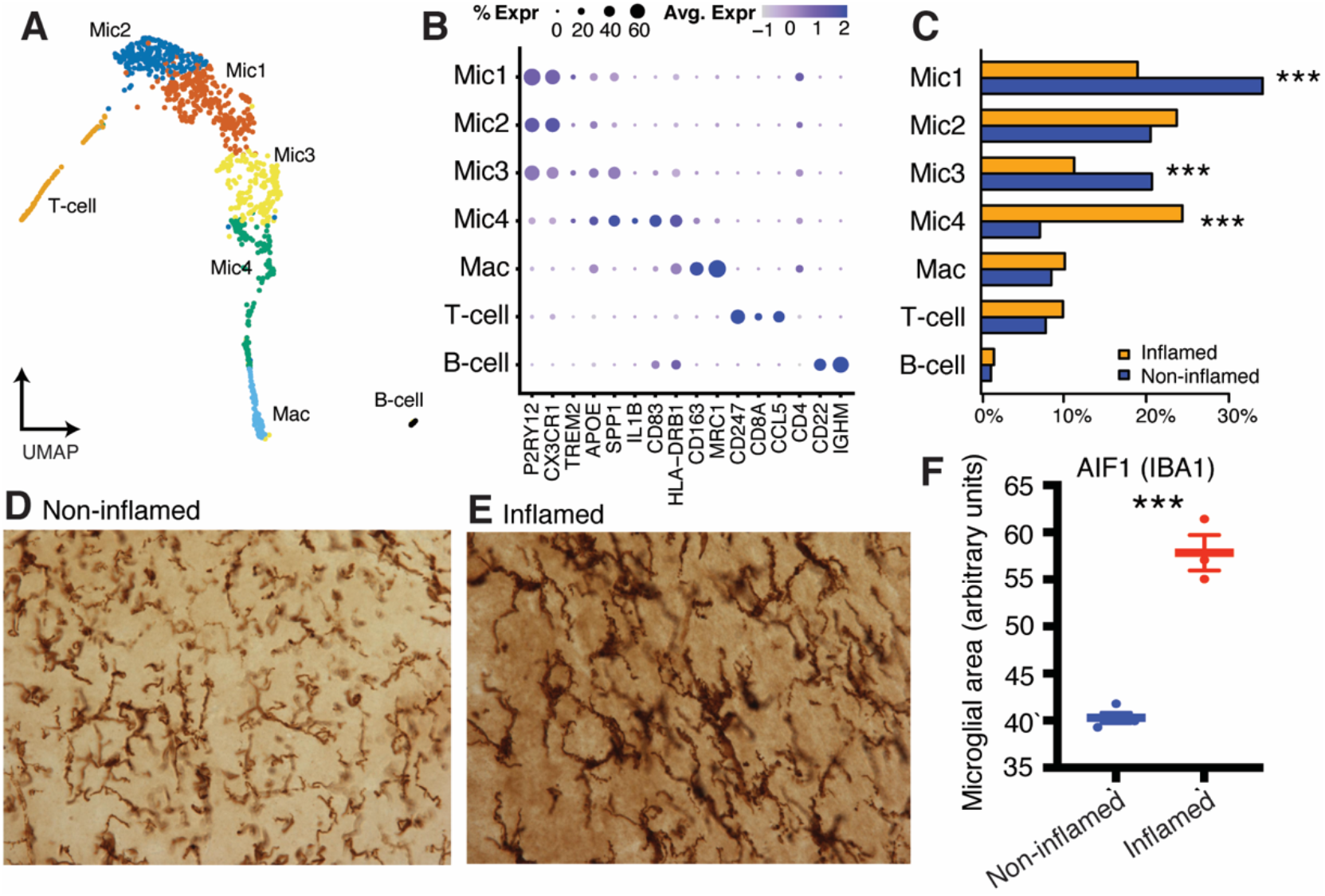
Transcriptional and morphological signatures of microglial reactivity in the cerebellum of children who died while experiencing inflammation. A. Sub-clustering of immune cells. B. Selected marker genes for immune cell subtypes. C. Differential abundance of microglial subtypes in inflamed vs. non-inflamed cerebelli. D,E. Representative images from immunohistochemical staining of microglia (IBA1, 20x) from a 2-year-old female (noninflamed) and 3-year-old female (inflamed). F. Microglial area (IBA1+) is increased in the cerebellum of inflamed vs. non-inflamed donors. *** *P* < 0.001.

Next, we calculated the effects of inflammation on differential gene expression in each cerebellar cell type. We identified a total of 5,710 cell type-specific gene expression changes (False Discovery Rate < 0.01, log_2_(fold change) > 0.5; Fig. 3A; Table S7). Significant gene expression changes were identified in all cell types, but by far the largest number of differentially expressed genes were detected in Golgi neurons followed by Purkinje neurons (Fig. 3B-C). Plots of top differentially expressed genes in these cell types confirmed that these effects were remarkably consistent across donors (Fig. S1). Both up- and down-regulated genes in these cell types were enriched for functional categories related to neurodevelopment and synapses, including strong enrichment for gene networks regulated by neuronal RNA-binding proteins such as FMRP, RBFOX, and CELF4 (Fig. 3D; Table S8), suggesting specific subtypes of cerebellar neurons may be selectively vulnerable to brain inflammation during childhood.

**Figure 3.**
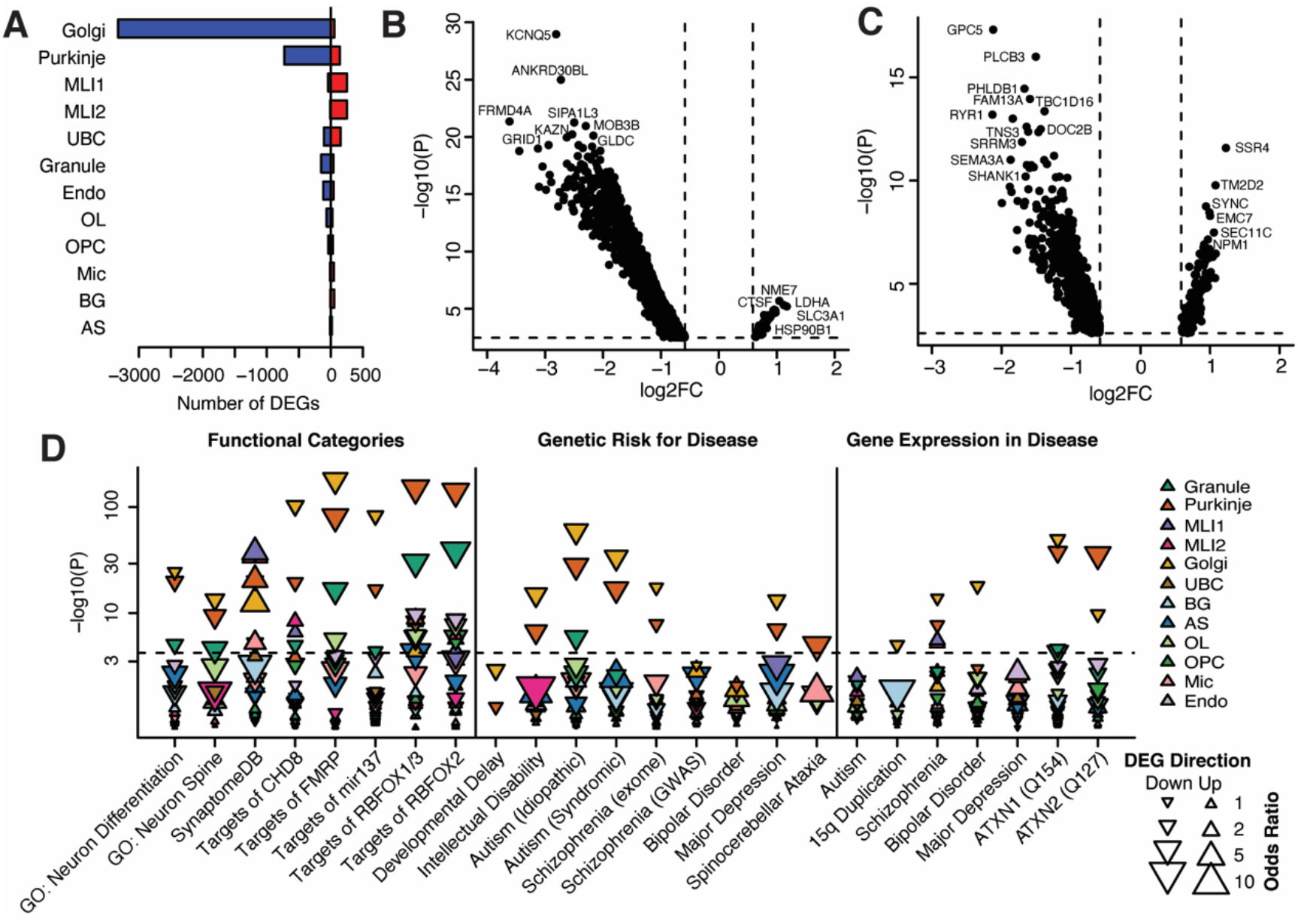
Cerebellar cell type-specific transcriptional consequences of inflammation. A. Counts of cell type-specific differentially expressed genes, FDR < 0.01, log2FC > 0.5). B,C. Volcano plots indicating fold-change and p-values for differential gene expression in Golgi neurons (B) and Purkinje neurons (C). D. Enrichments of differentially expressed genes in curated gene sets related to neurodevelopmental disorders. Down- and up-pointing arrowheads indicate enrichments among down-regulated and up-regulated transcripts, respectively. The y-axis is plotted on a log-scale to improve legibility.

As brain inflammation is both a risk factor and sub-phenotype in many psychiatric and neurological disorders (*1*), we tested whether genes influenced by inflammation are also implicated in neurological disorders. Down-regulated genes, primarily those in Golgi neurons and Purkinje neurons, were enriched for genes previously implicated in risk for intellectual disability, autism spectrum disorders, schizophrenia, and cerebellar ataxia (*31–34*) (Fig. 3D; Table S8). Down-regulated genes in these same cell types were also enriched for those differentially expressed in the post-mortem cerebellum of humans with schizophrenia or bipolar disorder (*35*), and in two knock-in mouse models of spinocerebellar ataxias (*Atxn1* Q154 and *Atxn2* Q127)(*36, 37*). Therefore, inflammation results in the down-regulation of many synaptic genes that are implicated in neurodevelopmental disorders and implicates specific subtypes of neurons.

### Transcriptional consequences of inflammation correspond to blunted maturation in Purkinje and Golgi neurons

In rats, inflammatory insults during the postnatal critical period result in a blunting of Purkinje neuron dendritic growth (*23*). We therefore calculated the effects of age on cerebellar gene expression in non-inflamed, 1-5-year-old donors and characterized the relationship between inflammation-induced and development-related changes. We detected substantial gene expression changes primarily in two neuronal subtypes, Purkinje neurons and Golgi neurons (Fig. 4A; Table S9). The majority of differentially expressed genes were down-regulated in cells from older (e.g., 3-5-year-old) donors, compared to 1- and 2-year-olds, consistent with our previous observation that the PGE2-E2 pathway is not engaged by inflammation in children one year or younger (*26*). Functional annotation suggests that these gene expression changes primarily involve decreased expression of genes involved in the formation of new synapses – e.g., the synaptic scaffolding genes *SHANK1* and *SHANK2,* which were down-regulated with age in Purkinje neurons – accompanied by a more subtle up-regulation of genes expressed at mature synapses. The maturation of Purkinje neurons and Golgi neurons was also associated with increased expression of genes related to energy metabolism and new protein synthesis, possibly reflecting the increased energetic demands of mature neurons.

**Figure 4.**
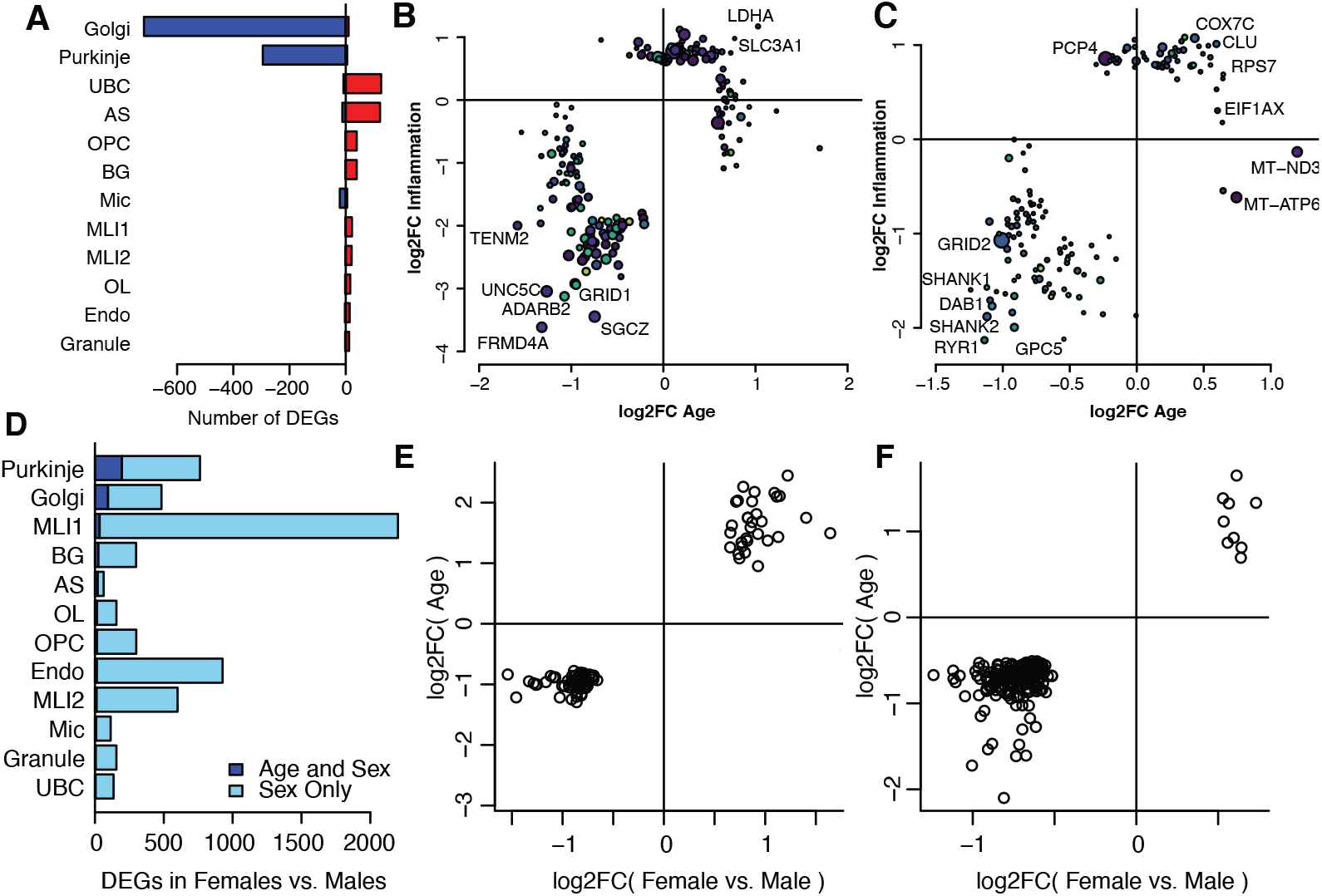
Correlated effects of age and inflammation in cerebellar cell types. A. Counts of genes with significant effects of age in 1- to 5-year-old non-inflamed donors (FDR < 0.01, log2(fold change) > 0.5). B,C. Correlated effects of age and inflammation in Golgi cells (B) and Purkinje cells (C). D. Counts of genes with significant sex differences in 1- to 5-year-old non-inflamed donors (FDR < 0.01, log2(fold change) > 0.5). Dark blue shading indicates the number of sex-different genes that were also influenced by age. E,F. In Golgi neurons (E) and Purkinje neurons (F), the directions of sex differences and the effects of age were positively correlated, such that at an equivalent age the transcriptomes in females were more mature than those of males.

We found strongly correlated transcriptional effects of inflammation and age both in Purkinje neurons and in Golgi neurons. In Golgi neurons, 388 of the 728 age-related genes were also differentially expressed in inflamed vs. non-inflamed donors, and the fold changes across all age- or inflammation-related genes were correlated (r = 0.56, *P* = 5.7e-224; Fig. 4B). Similarly, in Purkinje neurons, 81 of the 300 age-related genes were also differentially expressed with inflammation, and the effects were correlated (r = 0.66, *P* = 1.5e-129; Fig. 4C). These results suggest that the transcriptomic states for specific subtypes of cerebellar neurons undergo substantial changes during early childhood. Inflammation-related changes in these neurons appear to correspond to an acceleration of this maturation process.

The human cerebellum develops faster in girls than boys, and in animal models early life inflammation localized to the cerebellum has enduring deleterious consequences for males but not females (*23*). Sex differences in gene expression were detectable in all cerebellar cell types (Fig. 4D; Table S10). Among genes that were influenced by both sex and maturation, we found that the direction of these effects were correlated (across all cell types, Spearman’s rho = 0.36, *P* = 4.3e-15). Specifically, at an equivalent age, the transcriptomes of female donors appeared to be more mature than the transcriptomes of male donors. These effects were most strongly apparent in the same neuronal subtypes for which we observed substantial postnatal maturation, Golgi neurons and Purkinje neurons (Fig. 4E,F). We note, however, that the effects of inflammation, age, and sex are not independent, as they were measured in the same individuals.

Differences in the morphology of Purkinje neurons have been described in neurodevelopmental disorders such as autism spectrum disorders and schizophrenia, with the most reproducible association involving a reduction in soma size (*17*), and well as Purkinje neuron number and density (*38*). We quantified the size of Purkinje neuron soma from seven donors via immunostaining with calbindin (CALB1). There was no association of soma size either with age (r=0.2, *P* > 0.1) or between inflamed vs. non-inflamed donors (n=3-4 per group, Fig. S2). These results indicate that gene expression changes in Purkinje neurons from inflamed donors occur in the absence of dramatic, immediate changes in Purkinje neuron morphology. The lack of overt morphological differences argues against pre-existing pathological differences in the inflamed donors, none of whom had received a diagnosis with autism or a related neurodevelopmental disorder prior to death. Moreover, it is important to note that these children did not survive their inflammation. Thus, the enduring consequences for Purkinje morphology and survival are unknown. Gene expression changes occurring at the time of the inflammatory episode may set the stage for later cellular dysfunction (e.g., reduced synaptic connectivity and energy metabolism) that manifest as more frank differences in soma size after many years.

### Single-cell chromatin accessibility atlas for the human cerebellum

To gain insight into gene regulation in the cerebellum, we performed single-nucleus Assay for Transposase Accessible Chromatin (snATAC-seq) in the post-mortem cerebellum of one adult donor and one non-inflamed two-year-old child. We obtained high-quality chromatin accessibility profiles for 8.735 cells, which clustered to seven major cell types (Fig. 5A-C). Peak-calling within each cell type revealed a total of 135,773 accessible chromatin peaks, 8,857 of which were differentially accessible across cell types (FDR < 0.05, log2FC > 0.25; Table S11). Motif enrichment analysis identified sequence motifs enriched at the peaks specific to each cell type, including many corresponding to known cell type-specific regulatory factors (Table S12). Co-accessibility analysis identified 146,816 connections among these regions, predicting interactions between distal regulatory elements and the promoters they may regulate (Table S13).

**Figure 5.**
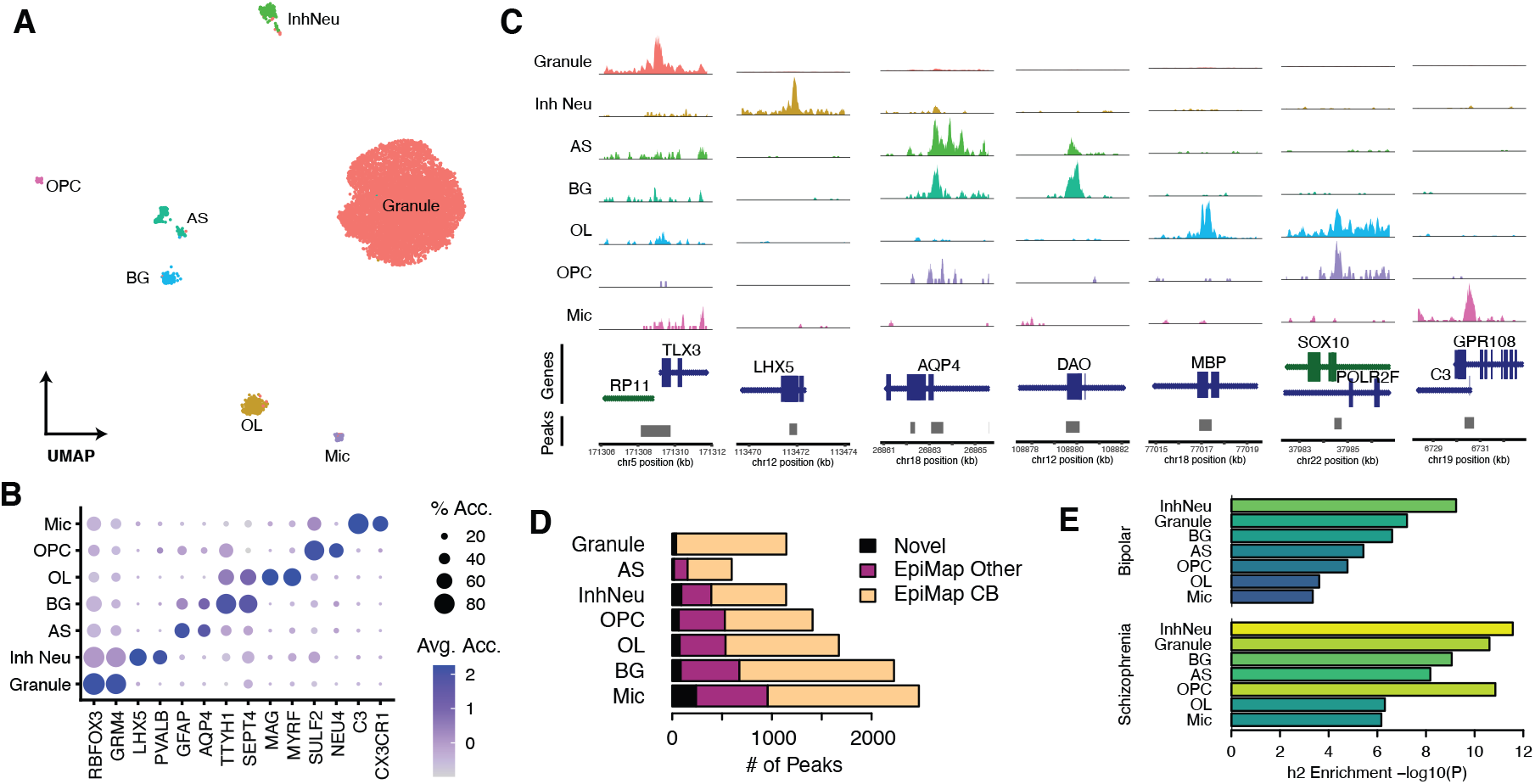
Single-cell chromatin accessibility atlas for the human cerebellum. A. Chromatin accessibility distinguishes major cerebellar cell types. B. Gene activity scores for selected cell type marker genes. B. Chromatin accessibility patterns for selected cell typespecific peak regions. D. Proportion of cell type-specific peaks overlapping known enhancers and promoters from EpiMap. E. Enrichment of cell type-specific accessible chromatin peaks in genomic regions associated with risk for psychiatric disorders.

We validated these candidate cis-regulatory elements via comparisons to annotations from EpiMap (*39*), 90,533 (67%) and 23,223 (17%) of the snATAC-seq peak regions overlap regions defined as enhancers or promoters, respectively, in any EpiMap sample. snATAC-seq revealed many regulatory regions in rare cell types that were missed by epigenomic profiling of bulk cerebellar tissue. Specifically, while 97% of the granule neuron-specific peaks in our dataset overlapped promoters and enhancers that were previously known to be active in the cerebellum based on EpiMap analyses of bulk cerebellar tissue, this was true for only 64% of peaks specific to other cell types (Fig. 5D).

Common genetic variants associated with risk for neuropsychiatric disorders are enriched at cis-regulatory elements active in the human brain. We tested whether heritable risk for two such disorders, schizophrenia and bipolar disorder, was enriched in the regulatory elements of specific cerebellar cell types utilizing stratified linkage disequilibrium score regression (*40*). Peak regions accessible in inhibitory neurons were the most strongly enriched for heritability in both disorders (Fig. 5E). This result supports the specific relevance of Purkinje neurons and other inhibitory neuron subtypes to neuropsychiatric disorders.

### An integrated model for the pre- and post-natal development of Purkinje cells and the effects of inflammation

Next, we sought to reconstruct the regulatory mechanisms underlying the maturation of human Purkinje neurons and the effects of inflammation. For this purpose, we integrated our snRNA-seq and snATAC-seq of Purkinje neurons with published snRNA-seq of fetal Purkinje neurons at 9-20 post-conception weeks (*27*). As a starting point for this analysis, we computed a gene co-expression network from the combined snRNA-seq data (n=1,531 Purkinje neurons), revealing 22 modules of co-expressed genes (Table S14). Seven of these modules were differentially expressed across maturation (Fig. 6A), with peak expression in early- to mid-gestation (M7, M12), late gestation (M14), one-year-olds (M2, M20), 2-5-year-olds (M15) and adults (M18). We also found bimodal expression of these modules in *ALDOC+* vs. *ALDOC-* sub-populations. Early-expressed modules were more highly expressed in *ALDOC-* Purkinje neurons, whereas late-expressed modules were more highly expressed in *ALDOC+* Purkinje neurons. We confirmed similar developmental patterns for six of these seven modules in publicly available gene expression profiles of sorted Purkinje neurons from the mouse cerebellum (*41*), suggesting that these modules are reproducible and evolutionarily conserved (Fig. S3).

**Figure 6.**
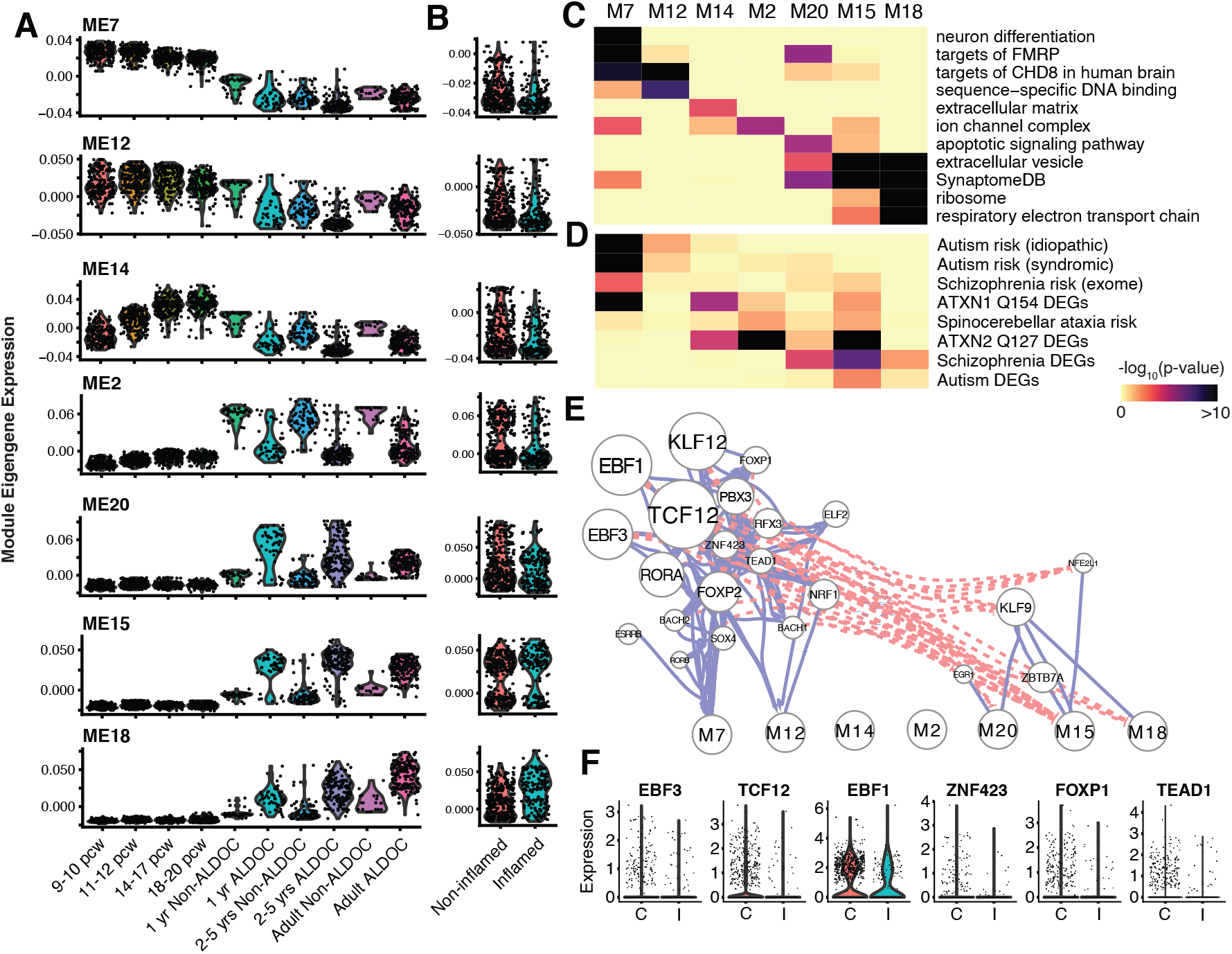
Gene regulatory network model for the maturation of the human cerebellum. A. Developmental- and subtype-specific expression of seven gene co-expression modules in human Purkinje cells, spanning from 9 post-conception weeks (PCW) to adulthood (47 years). B. Purkinje neurons maturation modules are differentially expressed in inflamed brains. C,D. Enrichments of gene co-expression modules for functional categories (C) and disease-associated genes (D). E. Integrative, gene regulatory network model predicts key transcription factors as regulators of Purkinje cell maturation. Solid, blue lines indicate activating interactions; dashed, red lines indicate inhibitory interactions. F. A subset of maturation-related TFs are down-regulated in Purkinje neurons from inflamed brains.

All seven of these developmentally regulated gene co-expression modules were further influenced by inflammation. The five modules that decreased across early childhood (M7, M12, M14, M2, and M20) were concordantly down-regulated in inflamed vs. non-inflamed brains. Both modules that increased across early childhood (M15, M18) were concordantly up-regulated in inflamed brains (Fig. 6B). That is, for each of these modules the effect of inflammation was equivalent to accelerated maturation. The effect sizes of inflammation were quite large, such that the transcriptional states of Purkinje neurons from an inflamed 1-2-year-old donor appeared almost equivalent to those from a non-inflamed 3-5-year-old donor.

We annotated the functional categories of genes at each stage of Purkinje neuron maturation (Fig. 6C; Table S15). Not surprisingly, modules with peak expression at prenatal timepoints were enriched for neurodevelopment processes (e.g., M2, “neuron differentiation”, *P* = 1.2e-35) and gene regulation (e.g., M12, “DNA binding”, *P* = 1.4e-10), whereas modules with peak expression postnatally primarily expressed genes for processes related to synapses (e.g., “SynaptomeDB”, *P* = 8.3e-7, 1.5e-27, and 1.2e-38 in M20, M15, and M18, respectively). M18, which reaches its full expression only in Purkinje neurons from adult donors, was further enriched for genes related to new protein synthesis and energy metabolism.

Genes expressed early vs. late in Purkinje neuron maturation also differed in their association with neurological disorders (Fig. 6D; Table S16). Genes for which inherited loss-of-function mutations are associated with risk for autism spectrum disorders or schizophrenia were enriched primarily in modules with peak expression levels prenatally (*32, 33, 35, 42*). By contrast, genes that are differentially expressed in post-mortem samples from each of these diseases were enriched in modules with peak expression postnatally, especially M15. Postnatal modules were also enriched for genes that are mutated in spinocerebellar ataxias. These results support a cerebellar neurodevelopmental etiology for autism and schizophrenia, and temporal distinctions between the expression patterns of genes associated with neurodevelopmental vs. neurodegenerative disorders of the cerebellum.

In order to reconstruct a gene regulatory network model for Purkinje neuron maturation we integrated our snRNA-seq and snATAC-seq datasets. We started by annotating sequence motifs in 14,853 accessible peaks proximal (<2kb) to the transcription start sites of Purkinje neuron-expressed genes, as well as 2,155 distal peaks predicted to interact with a proximal peak based on chromatin co-accessibility. We then fit a random forest regression model with GENIE3 to predict each gene’s expression as a function of the transcription factors (TFs) with putative binding sites in these regions. This resulted in a gene regulatory network model predicting the regulation of 7,837 target genes by 259 TFs, with a median of 6 regulators per target gene (Table S17). Of these interactions, 79% are predicted to be activating based on positive correlations between TFs and their predicted target genes. We next tested for enrichment of each TF’s predicted target genes within modules of co-expressed genes to predict key regulators at each stage of Purkinje neuron maturation (Fig. 6E; Table S18). Two distinct groups of regulators emerge. Predicted activators at the early phase of Purkinje neuron maturation (M7, M12) included established Purkinje neuron TFs such as *EBF1, RORA, FOXP2,* and *ZNF423,* as well as novel regulators such as *TCF12, SOX4,* and *PBX3.* These TFs typically are predicted to activate numerous other genes involved in neurodevelopment and gene regulation and to interact with one another through dense feed-forward loops. Predicted activators at the late stage of Purkinje neuron maturation included *KLF9, ZBTB7A,* and the activity-dependent TF *EGR1.* These TFs typically have fewer targets and primarily regulate effector genes such as synaptic components. The model also predicts inhibitory or non-linear interactions between many early and late regulators.

Finally, we evaluated gene regulatory networks to predict specific TFs that mediate inflammation’s effects on Purkinje neuron maturation. Our analyses highlight six TFs: *EBF3, TCF12, EBF1, ZNF423, FOXP1,* and *TEAD1.* Each of these TFs was predicted to regulate early phases of Purkinje neuron maturation, and both the TFs (Fig. 6F) and many of their predicted target genes (Table S19) were down-regulated in Purkinje neurons from inflamed brains. We used publicly available chromatin immunoprecipitation (ChIP-seq) data of FOXP1 in human neural progenitor cells (*43*) to validate our gene regulatory network model’s predictions. Consistent with our model, ChIP-seq-derived FOXP1 target genes were enriched for genes predicted to be activated by FOXP1 in our model (odds ratio = 2.0, *P* = 5.0e-3), genes in the M7 module (odds ratio = 2.4, *P* = 1.5e-8), and genes down-regulated in Purkinje cells from inflamed brains (odds ratio = 2.2, *P* = 5.1e-5). Plotting the developmental expression patterns for these TFs confirmed that each was highly expressed in prenatal Purkinje neurons and decreased within the first few years of postnatal life (Fig. S4). Our model therefore suggests that inflammation may blunt Purkinje neuron maturation by prematurely ending the postnatal expression of developmental gene regulatory programs in maturing Purkinje neurons.

## DISCUSSION

In recent years, new functions and increasing complexity of the cerebellum have been revealed in animal models by deep cellular phenotyping and opto- and chemogenetic interrogation (*44, 45*). However, roles of the cerebellum in human traits remain understudied compared to other brain regions. To our knowledge, this study represents the first comprehensive cell type-specific transcriptional and epigenetic characterization of the human cerebellum during childhood. It is also, to our knowledge, the first single-cell genomic study to characterize the effects of early-life inflammation in the brain. Use of a single-cell approach enabled us to dissect inflammation’s consequences in rare neuronal and glial cell types. Our results further establish the impact of early life inflammation on cerebellar development and its potential relevance to neurodevelopmental disorders.

Several limitations should be noted. There was considerable variation among samples in the sources of inflammation, ranging from meningitis to asthma. Nonetheless, we observed a remarkably high degree of congruency in transcriptional effects across subjects experiencing inflammation at the time of death. The metadata confirm no systematic differences between the two groups of subjects in variables such as post-mortem interval, time in freezer, race/ethnicity, age, or gender, and the detection of elevated COX-2 and/or TLR4 mRNA confirmed that there was inflammation in the brain. It is nonetheless possible that agonal state at the time of death is a common feature, resulting in systematic differences in oxidative stress in those subjects suffering inflammatory illness versus sudden death due to accident.

The specific cell types undergoing dramatic transcriptional changes in the contexts of maturation and inflammation included Golgi neurons, Purkinje neurons, and microglia. As all of these cell types are rare, many of these changes would be obscured in RNA-seq of bulk tissue, suggesting that previous studies may have underestimated the extent of neurodevelopmental disorder-related transcriptional vulnerability in the cerebellum (*42*). Notably, granule neurons, the most abundant cell in the cerebellum, were largely resistant to the impact of inflammation.

Golgi neurons were celebrated by their namesake, Camillo Golgi, for their large size, extensive arborization and localization at the interface of the cerebellar granule cell layer and molecular layer. After the initial anatomical characterization they largely fell into obscurity as attention was focused on the more beautiful and bountiful Purkinje neurons (*11*). Golgi neurons are predominantly GABAergic although a small percentage are exclusively glycinergic and others use both transmitters. Interest in Golgi cells re-emerged with modern electrophysiological and now transcriptomic approaches and has both confirmed the initial speculation of a role in creating networks and been expanded to include them as critical components of the recursive loops essential to temporal learning, plasticity, and transference of the acquisition of learned engrams to consolidation (*44*). They do not synapse with molecular layer interneurons but do form electrical synapses with each other, a feature that is presumed to contribute to even more refined temporal plasticity than previously believed (*44*). Our observation that Golgi neurons are particularly impacted by early life inflammation reveals the potential for deficits in multiple aspects of learning that may be subtle or context dependent and not evident as gross motor impairment.

Purkinje neurons were the next most impacted population by inflammation and as the sole output source from the cerebellum are of self-evident importance. Our gene co-expression network analyses identified seven development- and inflammation-associated modules, revealing the highly dynamic developmental trajectory of the cerebellum and potential for latent sensitive periods in early childhood. Revisionist views of the cerebellum emphasize multiple learning modes with the recent suggestion of a sensorimotor component that is independent of a second and distinct social emotional learning mode that relies on reward as a modulating influence (*44*). The discovery that sensorimotor learning is predominantly coded at ALDOC-Purkinje neurons (*13*) suggests the different modes may be coded at the level of different Purkinje neurons, but this remains to be determined.

Microglia are the innate immune cells of the brain and essential to healthy development but also central to dysregulation following neuroinflammation (*46–49*). Principal microglia functions include synaptic pruning (*50, 51*), promotion of synapse formation (*52*), and cell elimination (*53, 54*). Of the four microglia subtypes we identified with sub-clustering, one expressed pro-inflammatory markers and was highly enriched in the samples from subjects that perished with inflammation. Correspondingly, homeostatic subtypes were reduced in the same subjects. In addition, immunohistochemical staining revealed morphological features consistent with microglial reactivity, including increased microglial area. Artifacts in the transcriptomic profile of microglia following single cell or nucleus sequencing have been associated with enzymatic dissociation and *ex vivo* gene expression (*55*). While we cannot rule out that such an artifact exists in our data, all samples were processed identically and simultaneously, and there were no differences in post-mortem interval between inflamed vs. non-inflamed donors, suggesting that the shift in profile across inflammatory state is genuine.

The features of Golgi and Purkinje neurons that make them particularly vulnerable to inflammation induced dysregulation remain speculative. One shared feature is that these two cell types are the largest cells in the cerebellum. Large cell size may confer greater sensitivity to metabolic stress resulting in a higher number of differentially expressed genes. However, the differentially expressed genes identified were not necessarily indicative of cellular stress but instead more associated with neural development and synaptic processes. These enrichments point toward alternative explanations, such as the complex arborization and network engagement of both cell types. Indeed, it may be the morphological complexity of both Golgi and Purkinje neurons that necessitates their extended, postnatal maturation and makes them more vulnerable to inflammation, especially in comparison to the metabolically active but comparatively simple granule neurons. Importantly, inflammation in early childhood may also influence the maturation of neuronal subtypes in other brain regions, especially other brain regions with protracted postnatal maturation. Lastly, the goal of our study has been to elucidate the gene expression profiles of specific cells in the cerebellum as it develops and how that profile is modified by early life inflammation in order to understand how those modifications impact children who recover from illness. Characterizing the transcriptome in subjects that did not survive provides an imperfect but valuable proxy for attaining that goal.

Infections that occur specifically within the cerebellum -- as opposed to more general inflammatory episodes investigated here -- are rare in children and are usually diagnosed by acute ataxia in a previously healthy child (*56*). Viral infection is the most common cause and can progress from ataxia to cerebellitis, an even rarer inflammatory condition with a highly variable clinical presentation that ranges from listlessness, headaches and vomiting to death from compression of the brainstem. Given that recognition of the potential for inflammation within the cerebellum relies on severe motor impairment, headaches or neurological symptoms, it is likely that lesser degrees of damage from peripheral inflammation go undetected while nonetheless permanently altering the developmental trajectory. Moreover, reliance on symptoms such as ataxia and complaints of headaches will not detect inflammation in infants not yet walking and talking. Thus, there is considerable potential for undetected damage to the developing cerebellum as a consequence of viral or bacterial infections both within and outside the nervous system.

The relevance of early-life inflammation to neurodevelopmental and neuropsychiatric disorders is supported by the enrichments of inflammation-related gene expression changes for genes implicated in risk for these conditions. Previous studies demonstrated that Purkinje neurons are reduced in number and size in neuropsychiatric disorders, including autism spectrum disorders and schizophrenia (*57, 58*), with the most dramatic differences in the vermis, from which Purkinje neurons project to limbic regions, as well as in lateral regions of lobules VI and VII, which project to prefrontal cortex and contribute to cognition. Conditional knockout of several autism risk genes specifically in Purkinje neurons resulted in autistic-like behavioral changes (*59–63*), providing causal evidence for the involvement of these cells. Associations of Golgi neurons with neuropsychiatric disorders are largely unknown, suggesting a need for comparable studies to investigate these effects.

## MATERIALS AND METHODS

### Study design

The primary goal of this study was to assess cell type-specific transcriptional and epigenomic differences in the cerebellum of children who died while experiencing inflammation vs. non-inflamed controls. Post-mortem cerebellum tissue from human donors was studied using single-nucleus RNA sequencing single-nucleus ATAC sequencing on fresh frozen tissue, and immunohistochemistry on paraffin embedded fixed tissue.

### Human cerebellum samples

Post-mortem cerebellum tissue of 20 human donors was obtained from the University of Maryland Brain and Tissue Bank, the Human Brain Collection Core within the National Institute of Mental Health’s (NIMH) Division of Intramural Research Programs, and the Maryland Brain Collection at the Maryland Psychiatric Research Center (Table 1). Sample selection started from donors who were between 1 and 5 years old (UMB-BTB, NIH-HBCC) or >18-years-old (MBC) at the time of death, excluding donors who were previously diagnosed with any neurological or psychiatric disease. An investigator unaware of our experimental predictions and not involved in this study reviewed the limited medical records available and classified each donor as “inflamed” if subjects either: (1) had a cause of death indicated or exacerbated by infection, asthma, asphyxia, or inflammatory tumors, or (2) were treated with antibiotics or non-steroidal anti-inflammatory drugs around death. If none of these were present subjects were classified as “non-inflamed.” A subset of these donor brains were included in our previous study, in which we evaluated the expression of selected markers for inflammation. Tissue pH, time in freezer, post-mortem interval (PMI), and, in some cases, Aligent RNA integrity number (RIN) were recorded (*26*). All samples were expunged of any personally identifiable information prior to receipt. The study was reviewed by the University of Maryland Institutional Review Board and deemed “Not Human Subjects Research.”

### >Single-nucleus RNA sequencing

Nuclei were isolated from frozen brain tissue following a published protocol (*64*) with our slight modifications (https://dx.doi.org/10.17504/protocols.io.5jyl8973dv2w/v1). Briefly, frozen brain tissue was disaggregated by pipetting up and down in chilled extraction buffer consisting of Poly(1-vinylpyrrolidone-co-vinyl acetate) (Sigma #190845), 0.1% TritonX-100, and 1% bovine serum albumen (BSA) in dissociation buffer (DB; 82 mM Na_2_SO_4_, 30mM K_2_SO_4_, 10mM Glucose, 2.5mM MgCl_2_,10mM Hepes pH7.4, and RNase inhibitor [Protector, Roche]). Pellets containing nuclei were resuspended in DB and filtered with 70 um and 40 um strainers. To separate nuclei from debris, the nuclei suspension was mixed in a 25% (final) iodixanol layer over a 27% iodixanol cushion, followed by centrifugation (13000g x 20min, 4° C), then nuclei were washed twice in DB. Nuclei were counted using a MoxiGoII (Orflo) cytometer, and the concentration was adjusted to 365 nuclei/ul in phosphate buffered saline with 2% BSA. 17,000 nuclei were loaded in each well of a Chromium Controller (10x Genomics). Sequencing libraries were prepared using NextGEM 3’ Gene Expression reagents (10x Genomics), following manufacturer’s instructions and sequenced on HiSeq4000 (batches 1 and 3) and NovaSeq6000 (batch 2) sequencers.

### Single-nucleus ATAC sequencing

Cellular nuclei were isolated from frozen brain tissue following the Chromium Single Cell ATAC Demonstrated Protocol (CG000212 Rev C). Briefly, frozen samples were triturated in a disposable homogenizer (Biomasher, Takara) with 0.1X Lysis Buffer. Lysates were diluted with 0.5ml wash buffer and filtered with 70 um and 40 um strainers. Nuclei were centrifuged at 500g x5min, resuspended in diluted nuclei buffer at 3,000 nuclei/ul. Each sample was processed individually to this stage, then nuclei from a male and female donor (Table S1) were pooled to target 10.000 nuclei in total. Tn5 transposition and snATAC-seq library construction were performed using the 10x Genomics ATAC kit (v1), following manufacturer’s instructions, and sequenced on a NovaSeq6000 sequencer.

### Immunohistochemistry and morphological quantification

Samples of post-mortem cerebellum stored in 10% formalin were obtained from the University of Maryland School of Medicine Brain and Tissue Bank (https://www.medschool.umaryland.edu/btbank/) and sectioned by cryostat at 45um before mounting on silane-coated slides and washed in TBS (0.05M, pH = 7.6). Primary antibodies used were: mouse anti-calbindin (3-11 μg/ml; Sigma Cat# C9848, RRID: AB_476894); and rabbit anti-IBA1 (0.05-0.07 μg/ml, Wako Cat# 019-19741; RRID: AB_839504). Secondary antibodies used include the following: biotinylated goat anti-rabbit (3 μg/ml, Vector Cat#BA-1000, RRID: AB_2313606) and biotinylated horse anti-mouse (3 μg/ml, Vector Cat#BA-2000; RRID: AB_2313581). All incubations were performed at room temperature unless stated otherwise. To enhance immunolabeling of CALB1, antigen retrieval (0.01M sodium citrate, pH = 6.0 for 20 min at 95°C) was performed and slides were cooled to room temperature and incubated in a peroxide solution (50% histology-grade methanol/0.3% H2O2 in TBS) for 30 min. Slides were blocked with 5% normal goat serum (NGS) in TBS containing 0.4% Triton X-100 (TBST) for 1 h, followed by incubation in primary antibody solution containing 5% NGS in TBST overnight. The next day, slides were incubated with secondary antibody solution (2.5% NGS in TBST) for 1 h, incubated in ABC reagent (1:500 dilution; Vectastain Elite ABC Kit, Vector Laboratories) in TBST for 1 h, and visualized using DAB chromagen (0.05%3,3’diaminobenzidine, 0.006% hydrogen peroxide; Sigma-Aldrich D-5905). The DAB reaction was allowed to proceed until completion (~3-5 min). Sections on slides were dehydrated in a series of ascending ethanol, defatted in 100% xylene, and coverslipped with DPX mounting medium (VWR International Inc). Bright field images were captured on a Nikon Eclipse E600 using Neurolucida (v2020) software. Soma size of Purkinje neurons was measured at 20X by hand tracing and area quantified for 50-60 cells from 4-5 sections for each subject. The tissue was not of sufficient quality to allow for reliable quantification of cell number or density, but given the normal size a large decrease in cell number in inflamed individuals seems unlikely. Microglia morphology was assessed at 40X using ImageJ software after conversion to 8-bit gray scale and automatic thresholding, followed by measurement of area. The entire field of view was measured in four separate regions for each subject. In both cases data were averaged and used as the single measure per subject.

### snRNA-seq data processing, cell clustering, and annotation

Sequencing reads were processed to cell x gene read counts matrices using cellranger count v5 (10x Genomics, Pleasanton, CA), using the –include_introns flag and otherwise default parameters. In addition to standard quality control filters applied by CellRanger, we retained cells with at least 600 unique molecular identifier (UMI) counts. We used the Seurat v3 SCTransform pipeline (*65*) to integrate these cells across donors (samples from children and adults together) using 3,000 highly variable genes and 30 dimensions. We used these integrated counts for iterative cell clustering with Seurat. First, top-level clustering defined major cell types. Granule neurons were defined at this stage. We then re-integrated and sub-clustered interneurons, immune cells, and other nonneuronal cells (as three groups) to refine these cell populations. At each stage of clustering, we used 30 principal components with clustering resolution parameters derived manually to optimize cluster boundaries. We annotated the resulting cell clusters to known cerebellar cell types based on markers from a single-cell transcriptomic atlas of the mouse cerebellum. Clusters with mixed markers, corresponding to doublets and other low-quality cells, were dropped from downstream analyses.

### Differential gene expression

Effects of inflammation and age on gene expression were calculated for each cell type in which we identified at least 400 cells in samples from children. First, we reduced batch effects between cells from groups of samples processed on different days using ComBat-seq (*66*). Using edgeR, (*67*)we then fit a quasi-likelihood negative binomial generalized log-linear model to batch-adjusted counts, including main effects of group (inflamed vs. non-inflamed), age, and sex. For each cell type, differentially expressed genes associated with inflammation and age were defined as protein-coding transcripts detected in at least 20% of cells in either group, with p-values below a threshold corresponding to a 1% False Discovery Rate and log2(fold change) > 0.5. A small number of genes with >2-fold differences between male and female donors (primarily genes on sex chromosomes) had significant p-values in some analyses but are likely false positives due to imbalances in the number of cells from male vs. female donors and were therefore filtered from the final gene lists.

### Gene set enrichment analysis

We tested for over-representation of differentially expressed genes and gene co-expression modules in curated gene sets and Gene Ontology categories using Fisher’s exact tests. Curated gene sets were derived from Gene Ontology(*68*), SynaptomeDB, (*69*), Genovese et al. (2016)(*70*), the Deciphering Developmental Disorders consortium (*31*), SFARI Gene (*33*) SCHEMA (*32*) the Psychiatric Genomics Consortium (*71, 72*), Howard et al. (2019)(*73*), the Kyoto Encyclopedia of Genes and Genomes (*34*), Parikshak et al. (2016) (*42*), Chen et al. (2013)(*35*), and HDSigDB (*74*). Complete gene lists are shown in Table S19.

### snATAC-seq data processing, cell clustering, and annotation

Alignment of sequencing reads to the human genome (hg38) and initial cell and peak calling were performed with cellranger-atac v2. Cells from the two donors were de-multiplexed based on their genotypes using freemuxlet (*75*) then integrated with Signac (*76*) using reciprocal latent semantic indexing (LSI). Integrated counts were used for an initial round of clustering with Seurat to define major cell types. Next, we performed cell type-specific peak-calling to better define peak regions in rare cell types, using the MACS v2.1 (*77*) wrapper within Signac and the fragments assigned to each cell type. Read counts within these refined peak regions formed the basis for a second round of integration and clustering with Signac and Seurat. We selected high-quality cells with > 1000 and < 20,000 peak region fragments, > 15% of reads in peaks, < 5% of reads in blacklist regions, nucleosome signal < 4, and transcription start site enrichment > 2. Cells from the two donors were integrated using 29 reciprocal LSI dimensions, followed by clustering with 29 integrated dimensions, smart local moving (SLM) modularity optimization, resolution = 0.3, and otherwise default parameters. Cell clusters were assigned to cerebellar cell types based on accessibility within gene bodies (i.e., gene activity scores). Accessible peaks within each cell type were defined as those with fragments in at least 4% of cells of that type. Cell type-specific peaks were defined as those with a statistically significant enrichment of fragments (adjusted p-value < 0.05), based on Wilcoxon signed-rank tests implemented with FindMarkers(). Peaks were annotated based on overlap with UCSC hg38 knownGene gene models using the TxDb.Hsapiens.UCSC.hg38.knownGene and ChIPseeker R packages (*78*). Peaks were annotated to known promoters and enhancers using the GenomicRanges R package and annotations from EpiMap, (*39*) downloaded from http://compbio.mit.edu/epimap/ on April 29, 2022 and lifted to hg38, as necessary: enhancer masterlist (ENH_masterlist_locations.hg38.bed), promoter masterlist (PROM_masterlist_locations.hg38.bed), and sample-specific lists of promoters and enhancers for three cerebellar cortex samples, BSS00206, BSS00207, and BSS00201. Chromatin interactions were predicted using Cicero (*79*), following down-sampling to 500 cells per cell type to reduce bias toward granule neurons, using UMAP coordinates from Seurat, a maximum distance (“window”) of 250 kb, a distance constraint of 100 kb, and a co-accessibility threshold of 0.5.

### Heritability enrichment analysis

We used cell type-specific LD score regression (LDSC) to test for the enrichment of disease-associated SNPs in each cell type’s accessible chromatin regions (*80*). The merged set of accessible peak regions from all cell types was used as a baseline to control for non-cell type-specific associations of disease risk in accessible chromatin regions. Genome-wide association study summary statistics for schizophrenia (*81*) and bipolar disorder (*82*) were downloaded from the Psychiatric Genomics Consortium website (https://www.med.unc.edu/pgc/download-results/) on April 6, 2022.

### Gene co-expression network

We merged read counts in Purkinje neurons from our snRNA-seq datasets with published snRNA-seq of Purkinje neurons in the fetal brain (*27*)Read counts and cell annotations from fetal samples were downloaded from the UCSC Cell Browser (https://cbl-dev.cells.ucsc.edu, version 1). Prior to network reconstruction, read counts were normalized in six steps: (i) genes with read counts in < 3% of cells were filtered; (ii) matrices were downsampled to ≤ 100 cells per donor; (iii) ComBat-seq was applied to reduce batch effects, with group = inflammation and covar_mod = log10(age); (iv) knn-smoothing (*83*) (k=3, d=30) was applied to reduce dropouts; (v) smoothed counts were normalized to counts per million (Seurat NormalizeData function); then (vi) mean-centered and scaled to unit variance. Next, modules of co-expressed genes were identified by k-means clustering, using the kmeans function from the stats package in R with k=25, iter-max=10000, nstart=10, and algorithm=“Lloyd”. Genes with weak correlations to module centroids (Pearson’s r < 0.2) were removed. Modules with very similar patterns of expression were merged via average-linkage hierarchical clustering of cluster centroids, using 1 – Pearson’s r as a distance metric and a cut height of 0.1. Associations of modules with age and inflammation were calculated based on Fisher’s exact test enrichments for differentially expressed genes. For plotting purposes, modules were summarized by their first principal components, calculated using the moduleEigengenes() function from the WGCNA R package (*84*). For validation, we studied RNA-seq of sorted Purkinje neurons from the developing mouse cerebellu (*41*). Read counts and sample metadata were downloaded from the Gene Expression Omnibus (GSE86824). We plotted the average expression of log2-normalized counts per million for the mouse orthologs of the genes in each module.

### Integrated gene regulatory network

We reconstructed a gene regulatory network (GRN) model for the regulation of Purkinje neuron gene expression by transcription factors, utilizing GENIE3 and constraining the algorithm to consider only those transcription factors with putative binding sites in each gene’s accessible promoters and enhancers. The general framework is to fit a random forest regression model to predict the expression of each target gene by the combined expression patterns of transcription factors (TFs). GENIE3 has repeatedly been shown to outperform other methods that have been proposed for this task. In previous work, we and others have demonstrated that constraining the search space by the locations of sequence motifs improves the precision and recall for predicting direct target genes. Our implementation had three inputs: (i) Putative transcription factor binding sites within each accessible peak region in inhibitory neurons, derived via a motif scan with the Signac AddMotifs() function, point-weight matrices from the JASPAR2020 R package, and motif-to-TF mappings from JASPAR and MotifDb; (ii) peak-to-gene mappings, including proximal peaks < 2 kb from a transcription start site and distal peaks predicted by Cicero to interact with a proximal peak; (iii) centered and scaled gene expression data from Purkinje neurons, as described in the previous section. Inputs (i) and (ii) are combined to produce a “base” GRN describing the set of TFs with predicted binding sites within each gene’s putative regulatory elements. These TFs were provided as input to GENIE3. GENIE3 ranks the TF-to-gene interactions by an importance score derived from the random forest regression. We selected the top 50,000 TF-to-gene interactions from the model, corresponding to a median of five predicted regulators per gene, and we assigned each TF-to-gene interaction as activating or inhibitory based on the sign of the Pearson correlation between the pair of genes. TF-to-TF interactions are a special class of interaction that can be obtained directly from the model. Associations of TFs with gene co-expression modules and inflammation were predicted based on Fisher’s exact test over-representation for each TF’s predicted target genes (activating and inhibitory targets separately). We validated our model’s predictions for *FOXP1* using experimentally-derived target genes from FOXP1 ChIP-seq in human neural progenitor cells (Supplementary Table 1 from (*43*)).

### Statistical analysis

Statistical analyses for single-cell genomics experiments are described in the preceding sub-sections. For immunohistochemistry experiments, statistical comparisons between groups were assessed with t-tests.

## List of Supplementary Materials

Fig. S1 to S4

Data files S1 to S19 (Excel files)

## Acknowledgments

We thank the staff of the University of Maryland Brain and Tissue Bank, the Human Brain Collection Core within the National Institute of Mental Health’s (NIMH) Division of Intramural Research Programs, and the Maryland Brain Collection at the Maryland Psychiatric Research Center for their assistance in procuring post-mortem brain tissue and associated metadata. We thank Sheryl Arambula, PhD for sample preparation. Sequencing was performed by the Genomics Resource Center within the Institute for Genome Sciences at the University of Maryland School of Medicine.

## Author contributions

Conceptualization: SAA, MMM

Methodology: SAA, MCG, CC

Investigation: SAA, MCG, BRH, EM, MMM

Visualization: SAA, MMM

Funding acquisition: SAA, MMM

Project administration: SAA, MMM

Supervision: SAA, MMM

Writing – original draft: SAA, MMM

Writing – review & editing: SAA, MCG, BRH, EM, CC, MMM

## Data and materials availability

Raw and processed single-cell genomic data have been deposited in the Neuroscience Multi-Omic Archive. A web-based tool for the visualization of the human cerebellum snRNA-seq is available at NeMO Analytics (https://nemoanalytics.org/p?s=522872b1). Custom R scripts for the reconstruction of gene regulatory networks are available on GitHub (https://github.com/seth-ament/cbdev). All other data are available in the main text or the supplementary materials.

## SUPPLEMENTARY MATERIALS

**Figure S1.**
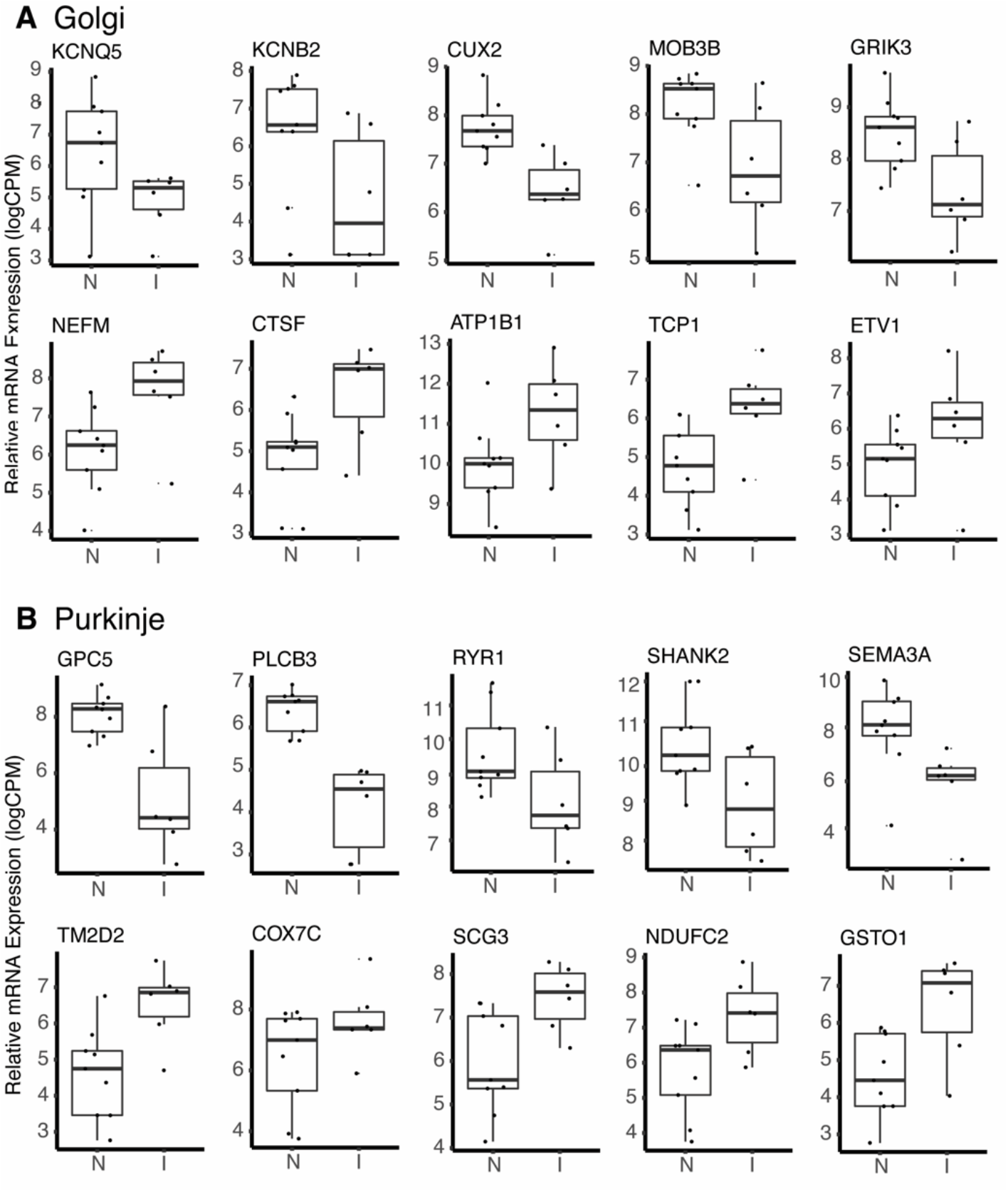
Expression patterns of top differentially expressed genes in Golgi neurons and Purkinje neurons. Plots indicate pseudobulk log-transformed counts per million, with dots representing the aggregate expression level in each donor brain. Note that cells from two donors are not shown in the pseudobulk plots due to a relatively low number of cells being captured from those donors. Genes shown were selected from among the top 30 up- and down-regulated genes with the lowest p-values in each cell type. A. Golgi neurons. B. Purkinje neurons.

**Figure S2.**
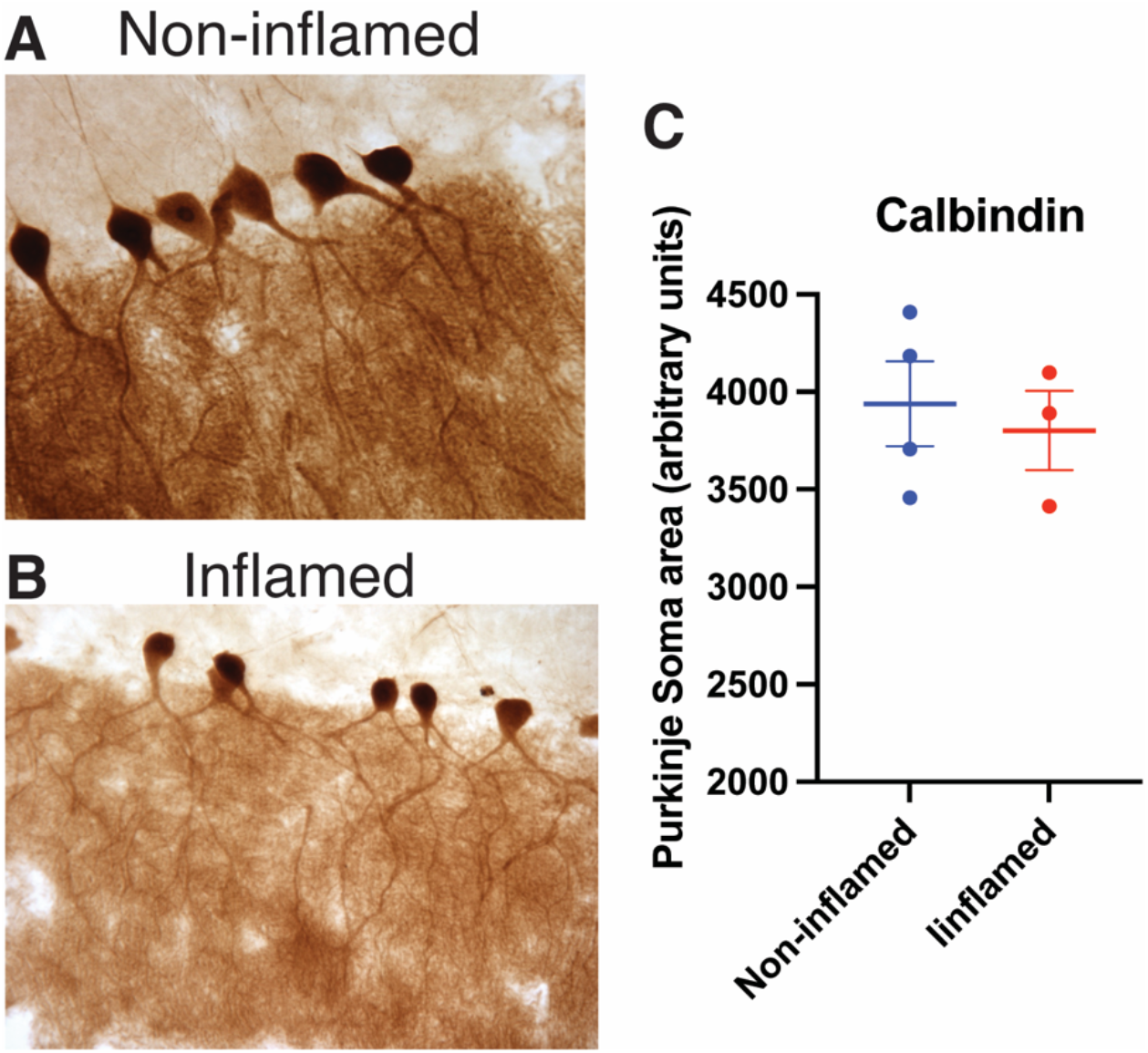
Immunostaining of Purkinje neurons in the cerebellum of inflamed vs. noninflamed donors. A,B. Represent images of Calbindin (CALB1) immunoreactive cells (20x objective). C. Quantification of Purkinje neuron soma size (*P* > 0.1).

**Figure S3.**
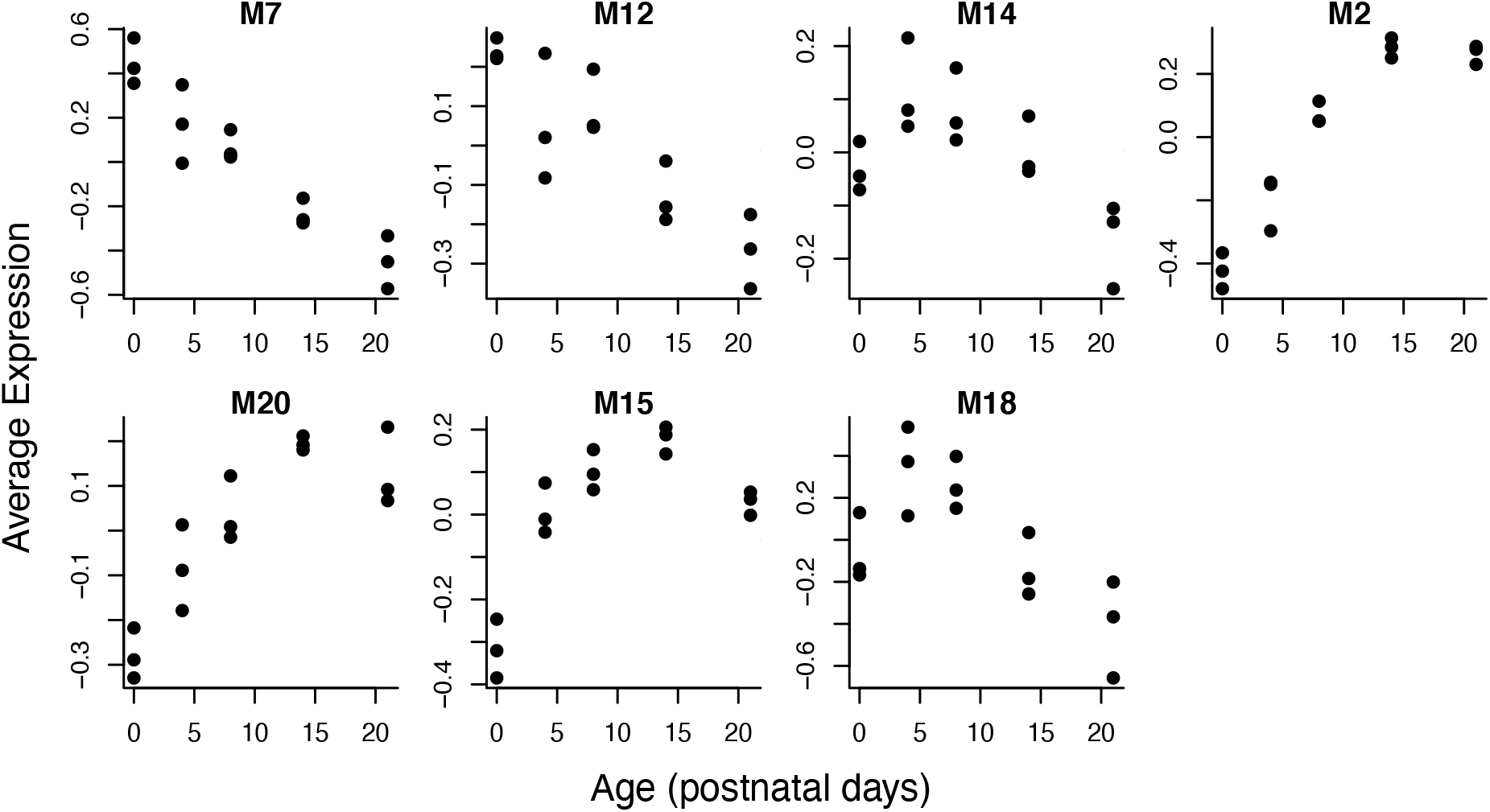
Developmental dynamics of human Purkinje neuron gene co-expression modules during the maturation of Purkinje neurons in mice. Mouse data are from GSE86824. Points indicate the average expression of log2-normalized counts per million for the mouse orthologs of the genes in each module.

**Figure S4.**
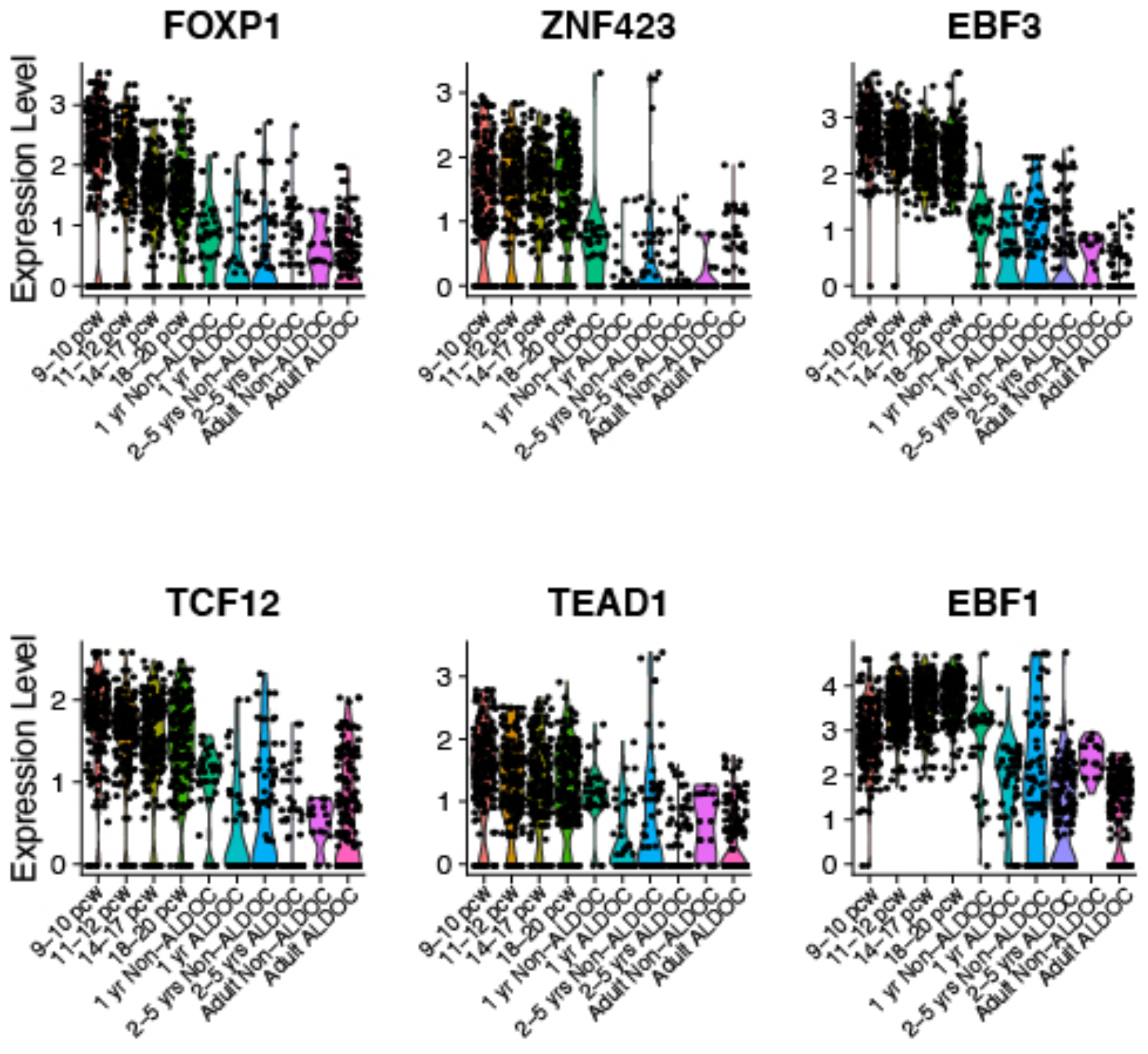
Developmental dynamics of key developmental and inflammation-related transcription factors in Purkinje neurons. Y-axis indicates normalized expression levels for each gene in snRNA-seq of Purkinje neuron sub-types at the indicated age ranges.

**Tables S1 to S19 (Excel files)**

**Table S1. Sample characteristics**

**Table S2. Cell type marker genes in snRNA-seq of cerebellar cortex samples from children**

**Table S3. Cell type marker genes in snRNA-seq of cerebellar cortex samples from adults**

**Table S4. Differential gene expression among molecular layer interneuron subtypes**

**Table S5. Differential gene expression among Purkinje neuron subtypes**

**Table S6. Differential gene expression among immune cell subtypes**

**Table S7. Differential gene expression in inflamed vs. non-inflamed donor brains**

**Table S8. Enrichment of inflammation DEGs for curated gene sets**

**Table S9. Differential gene expression effects of age**

**Table S10. Differential gene expression effects of age**

**Table S11. Differentially accessible chromatin peaks in cerebellar cell types**

**Table S12. Sequence motifs enriched in differentially accessible chromatin peaks**

**Table S13. Interactions among chromatin peaks predicted by co-accessibility**

**Table S14. Gene co-expression modules in maturing Purkinje neurons**

**Table S15. Enrichments of gene co-expression modules within functional categories**

**Table S16. Enrichments of gene co-expression modules within curated disease-related gene sets**

**Table S17. Predicted transcription factor-target gene interactions in human Purkinje neurons**

**Table S18. Enrichments of gene co-expression modules for the target genes of transcription factors**

**Table S19. Enrichment of inflammation DEGs for the target genes of transcription factors**

